# Multimodal Analyses of Stable Vitiligo Skin Identify Tissue Interactions That Control Disease Persistence

**DOI:** 10.1101/2021.12.03.470971

**Authors:** Jessica Shiu, Lihua Zhang, Griffin Lentsch, Jessica L Flesher, Suoqin Jin, Christopher Polleys, Seong Jin Jo, Craig Mizzoni, Pezhman Mobasher, Jasmine Kwan, Francisca Ruis-Diaz, Bruce J Tromberg, Irene Georgakoudi, Qing Nie, Mihaela Balu, Anand K Ganesan

## Abstract

Vitiligo is an autoimmune skin disease that is characterized by the progressive destruction of melanocytes by autoreactive CD8^+^ T cells. Melanocyte destruction in active vitiligo is mediated by CD8^+^ T cells but why white patches in stable disease persist is poorly understood. The interaction between immune cells, melanocytes, and keratinocytes *in situ* in human skin has been difficult to study due to the lack of proper tools. Here, we combine non-invasive multiphoton microscopy (MPM) imaging and single-cell RNA sequencing (scRNA-seq) to identify distinct subpopulations of keratinocytes in lesional skin of stable vitiligo patients. We show that these keratinocytes are enriched in lesional vitiligo skin and differ in metabolism, an observation corroborated by both MPM and scRNA-seq. Systematic investigation of cell-cell communication show that CXCL is the prominent signaling change in this small population of keratinocytes, which secrete CXCL9 and CXCL10 to create local inflammatory cytokine loops with T cells to drive stable vitiligo persistence. Pseudotemporal dynamics analyses predict an alternative keratinocyte differentiation trajectory that generates this new population of keratinocytes in vitiligo skin. In summary, we couple advanced imaging with transcriptomics and bioinformatics to discover cellcell communication networks and keratinocyte cell states that perpetuate inflammation and prevent repigmentation.

**One Sentence Summary:** Communication between keratinocytes, immune cells, and melanocytes maintain depigmented patches in stable vitiligo.

## INTRODUCTION

Vitiligo is an autoimmune skin disease characterized by the progressive destruction of melanocytes by autoreactive CD8^+^ T cells, resulting in disfiguring patches of white depigmented skin that cause significant psychological distress among patients (*1*). CD8^+^ T cells play an important role in the elimination of melanocytes and are increased in active vitiligo skin (*2*–*4*). However, in stable vitiligo lesions devoid of melanocytes, T cells are sparse and immune activation levels are low (*5*). This makes it unclear why white patches continue to persist in the absence of a robust inflammatory infiltrate.

Development of mouse models representative of human disease has provided important clues on the role of the adaptive immune system in vitiligo (*6*, *7*). Keratinocytes secrete CXCL9 and CXCL10 to attract and activate CXCR3^+^ CD8^+^ T cells (*8*) and these chemokines are present in the blister fluid of human vitiligo patients (*4*). However, the adoptive transfer of autoreactive CD8^+^ T cells in the mouse model cannot fully recapitulate the complex interactions between melanocytes, keratinocytes, and immune cells that occurs *in situ* in human skin-melanocytes are present in the epidermis in only select locations in mice (*9*) and the mouse epidermis is considerably thinner and lacks the stratification seen in human skin (*10*). To date, most translational studies in vitiligo are limited to examining cultured cells *in vitro* or immunohistochemistry of diseased tissue(*11*). It has been difficult to study how cell lineages collectively contribute to disease persistence secondary to the lack of tools to assess cellular heterogeneity *in vivo*.

Multiphoton microscopy (MPM) is a unique tool for this purpose and has broad applications in human skin (*12*–*19*). MPM is a noninvasive imaging technique capable of providing images with sub-micron resolution and label-free molecular contrast which can be used to characterize keratinocyte metabolism in human skin (*20*, *21*). This approach is based on the two-photon excited fluorescence (TPEF) signal detected from the reduced nicotinamide adenine dinucleotide (NADH), a co-enzyme in the keratinocyte cytoplasm that plays a central role in metabolism. We have validated this technique’s ability to assess cellular metabolism in normal skin under hypoxic conditions (*21*, *22*). Further addition of multiparametric analyses of NADH and flavin adenine dinucleotide (FAD) allows quantification of metabolic changes at a single cell resolution in cells and tissue(*23*).

In this study, we employ MPM for *in vivo* imaging of stable vitiligo lesions and assess keratinocyte metabolic state based on an imaging metric derived from a mitochondrial clustering analysis approach validated in previous studies (*21*, *22*). We then performed single-cell RNA sequencing (scRNA-seq) on patient-matched lesional and nonlesional tissue to identify keratinocyte subpopulations that express chemokines known to drive vitiligo pathogenesis. By applying CellChat, a tool that quantitatively infers and analyzes intercellular communication networks in scRNA-seq data, we demonstrate that stress keratinocytes communicate with adaptive immune cells via the CXCL9/10/CXCR3 axis to create local inflammatory loops that are active in stable vitiligo. Moreover, signaling between melanocyte and keratinocytes via the WNT pathway was altered in stable vitiligo lesions. By integrating non-invasive MPM, scRNAseq, and advanced bioinformatics, we infer communication networks between keratinocytes, melanocytes, and immune cells capable of preventing normal melanocyte repopulation.

## RESULTS

### MPM imaging of stable vitiligo skin *in vivo* demonstrate mitochondrial clustering changes

To look at epidermal changes using MPM in stable vitiligo, we utilized the MPTflex clinical microscope (see methods) to image twelve patients with lesions characterized by depigmented areas that have not grown in size for at least one year and did not exhibit active vitiligo features such as confetti-like depigmentation, koebnerization and trichome (table S1)(*24*). As expected, MPM images of nonlesional skin showed brighter fluorescence spots in the cellular cytoplasm, which represent aggregates of melanosomes, compared to lesional skin (fig. 1A) (*15*). To evaluate for metabolic changes in nonlesional and lesional vitiligo skin, we studied mitochondrial clustering which was previously validated in skin under normal and hypoxic conditions (*21*). Consistent with published data, nonlesional skin exhibited depth-dependent changes in mitochondrial clustering that reflects differences in metabolism (fig. 1A). In short, the basal and parabasal keratinocytes present a fragmented mitochondria phenotype characterized by high values of the mitochondrial clustering metric, β. As cell differentiation progresses from the basal to the higher epidermal layers and cells turn from glycolysis to oxidative phosphorylation for energy production, mitochondria fuse and create more extensive networks that correspond to lower clustering values, reaching their minima within the spinous layer (fig. 1A). Finally, toward the most terminally differentiated layer, as the granular keratinocytes enter an apoptotic state to create the stratum corneum, mitochondrial clustering values recover again, signifying a return to a more fissioned phenotype. In contrast, lesional depigmented skin from vitiligo patients showed an altered trend of mitochondrial clustering compared to nonlesional skin (fig. 1A), suggesting that the depth-dependent metabolic changes were lost. We calculated the mitochondrial clustering (β) median value and its variability across the epidermis of vitiligo and normal skin and found that these metrics are significantly different in vitiligo lesional and nonlesional skin (fig. 1C). Given that these changes were observed in the basal layer, we performed additional analysis to compare mitochondrial clustering between lesional and nonlesional basal keratinocytes. This analysis indicates a more heterogeneous distribution of mitochondrial clustering, β, values for lesional vitiligo vs non-lesional basal keratinocytes (fig. S1), yielding distributions with heterogeneity index values of 0.16 and 0.12 respectively. Noticeably, vitiligo basal keratinocytes exhibited an increase in the number of cells characterized by lower mitochondrial fragmentation levels and thus more networked mitochondria, consistent with enhanced oxidative phosphorylation (*21*–*23*).

**Fig 1.**
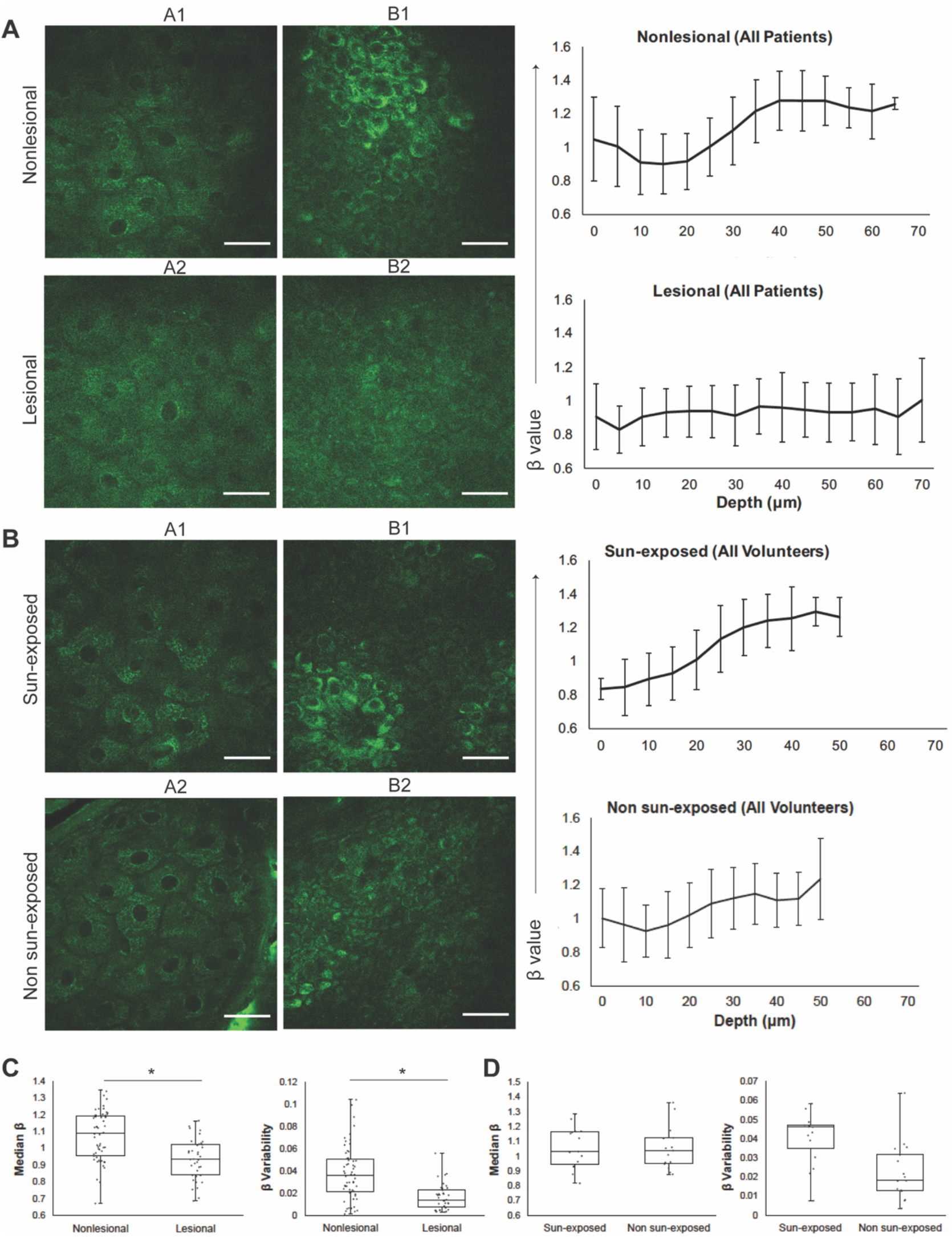
*In vivo* MPM images of vitiligo lesional and nonlesional skin showing metabolic changes with depth independent of exposure. **(A)** Representative en-face MPM images from the stratum granulosum in nonlesional (A1) and lesional skin (A2) and from the basal layer in nonlesional (B1) and lesional skin (B2) of one vitiligo patient. Average mitochondrial clustering (β) values based on z-stacks from all vitiligo patients as a function of depth for nonlesional (top right) and lesional (bottom right) skin are shown as spline fits. Error bars represent the standard deviation of the measurements for the images in all the z-stacks at each area. The labels A1, A2, B1, and B2 within the mitochondrial clustering panels represent the mitochondrial clustering values extracted from the panel’s respective labeled images. Scale bars are 20μm. **(B)** Representative en-face MPM images from the stratum granulosum in sun exposed (A1) and non sun-exposed skin (A2) and from the basal layer in sun exposed (B1) and non sun-exposed skin (B2) of healthy volunteers. **(C)** Distribution of the median β values (left) and β variability values (right) in nonlesional and lesional skin of vitiligo patients; each value corresponds to a z-stack of images acquired in nonlesional and lesional skin. * = t-test p-value < 0.05 **(D)** Distribution of the median β values (left) and β variability values (right) in sun-exposed and non sun-exposed skin of healthy volunteers; each value corresponds to a z-stack of images acquired in non sun-exposed and sun-exposed areas.

Since the fluorescence signals from all the skin fluorophores, including NADH, are collected on the same detection channel in the MPTflex, we sought to ensure the mitochondrial clustering measurements were not affected by contributions from fluorophores other than NADH. Melanin requires particular consideration since it is the main source of difference in appearance between vitiligo and normal skin. To ensure that melanin content was not affecting fluorescence signals, we measured mitochondrial clustering in five healthy volunteers. We controlled for melanin content by comparing sun exposed sites (dorsal forearm) and non-sun exposed sites (volar upper arm, which would have relatively less melanin). We found that depth-dependent β values showed similar trends in the epidermis (fig. 1B) regardless of sun-exposure status and the median β values and β variability values were not significantly different (fig. 1C). These results confirmed that mitochondrial clustering in basal and parabasal keratinocytes of lesional skin was altered compared to nonlesional skin. This was a result of changes to mitochondrial organization in vitiligo skin and was not a consequence of differences in melanin content.

### scRNA-seq reveals unique keratinocyte cell states enriched in vitiligo lesional skin

MPM imaging demonstrated that basal and parabasal keratinocytes in vitiligo lesions were metabolically altered, suggesting that keratinocyte cell states are different in vitiligo patients. To systematically examine the major keratinocyte cell state changes in vitiligo, we performed scRNA-seq on a separate group of patient-matched lesional and nonlesional suction blisters from seven patients using the 10x Genomics Chromium platform (fig. 2A). 1 set of samples (patient B) was excluded from further analyses due to the low viability of cells (Table S2). We performed read depth normalization and quality control (see Methods section, fig. S2), and obtained a total of 9254 cells of vitiligo lesional skin and 7928 cells of nonlesional skin for downstream analyses. We performed integration analysis of data from all patients using our recently developed approach scMC, which is designed to preserve biological signals while removing batch effects(*25*). Unsupervised clustering analysis identified 14 cell clusters (fig. 2B). Using the differentially expressed gene signatures, we were able to attribute clusters to their putative identities (fig. 2CD), including basal keratinocytes (high KRT15 and KRT5 expression), spinous keratinocytes (high KRT1 expression), granular keratinocytes (high FLG and LOR expression), cycling keratinocytes (high TOP2A expression), melanocytes (high PMEL expression), TC (T cell) (high CD3D expression) and DC (Dendritic cell) (high CD207 expression) (fig. 2B). The intermediate keratinocyte states, including basal to spinous transition and spinous to granular transition, were defined based on the hybrid expression of KRT15, KRT1 and KRT2. Notably, we identified two keratinocyte states that upregulate expression of keratins that are not normally expressed in the mature interfollicular epidermis and are associated with insults like wounding and UV injury (fig. 2B) (*26*, *27*). Stress 1 subpopulation was highly enriched for KRT6A while Stress 2 subpopulation expressed KRT6A at lower levels. They also expressed KRT16 and S100A8/9, alarmins associated with local inflammation that have been used as biomarkers for other inflammatory conditions(*28*). We term these populations “stress keratinocytes” as their transcriptional signature corresponds with injuries and inflammation. Interestingly, stress keratinocytes were only enriched in vitiligo lesional skin (fig. 2B). Detailed analysis of the two immune cell subpopulations TC and DC showed that they were distinguished from each other with clearly distinct gene signatures and biological processes (fig. S3). Cellular composition analysis showed that although different patients exhibited certain heterogeneity, cell clusters were common amongst patients (fig. 2e). Compared to nonlesional skin, vitiligo lesional skin showed dramatically increased presence of stress keratinocyte and to a lesser extent of DC, and a clear decrease of melanocytes (fig. 2e). Overall, the percentages of keratinocytes and melanocytes were decreased, and stress keratinocytes and immune cells were increased in vilitigo lesional skin (fig. 2f). Moreover, we analyzed keratinocytes from normal human skin using a previously published scRNA-seq dataset (fig. S4), but did not observe the expression of stress signature genes, suggesting that stress keratinocytes were uniquely enriched in vitiligo lesional skin. Integration analysis using a Seurat package produced similar cellular compositions, but did not preserve biological variation as well. In particular, stress keratinocytes were intermixed with other keratinocyte cell states and were in a spread distribution in the UMAP space (Uniform Manifold Approximation and Projection) (fig. S5). Collectively, these data provide the first general overview of the major changes in cellular compositions from nonlesional skin to vitiligo lesional skin.

**Fig 2.**
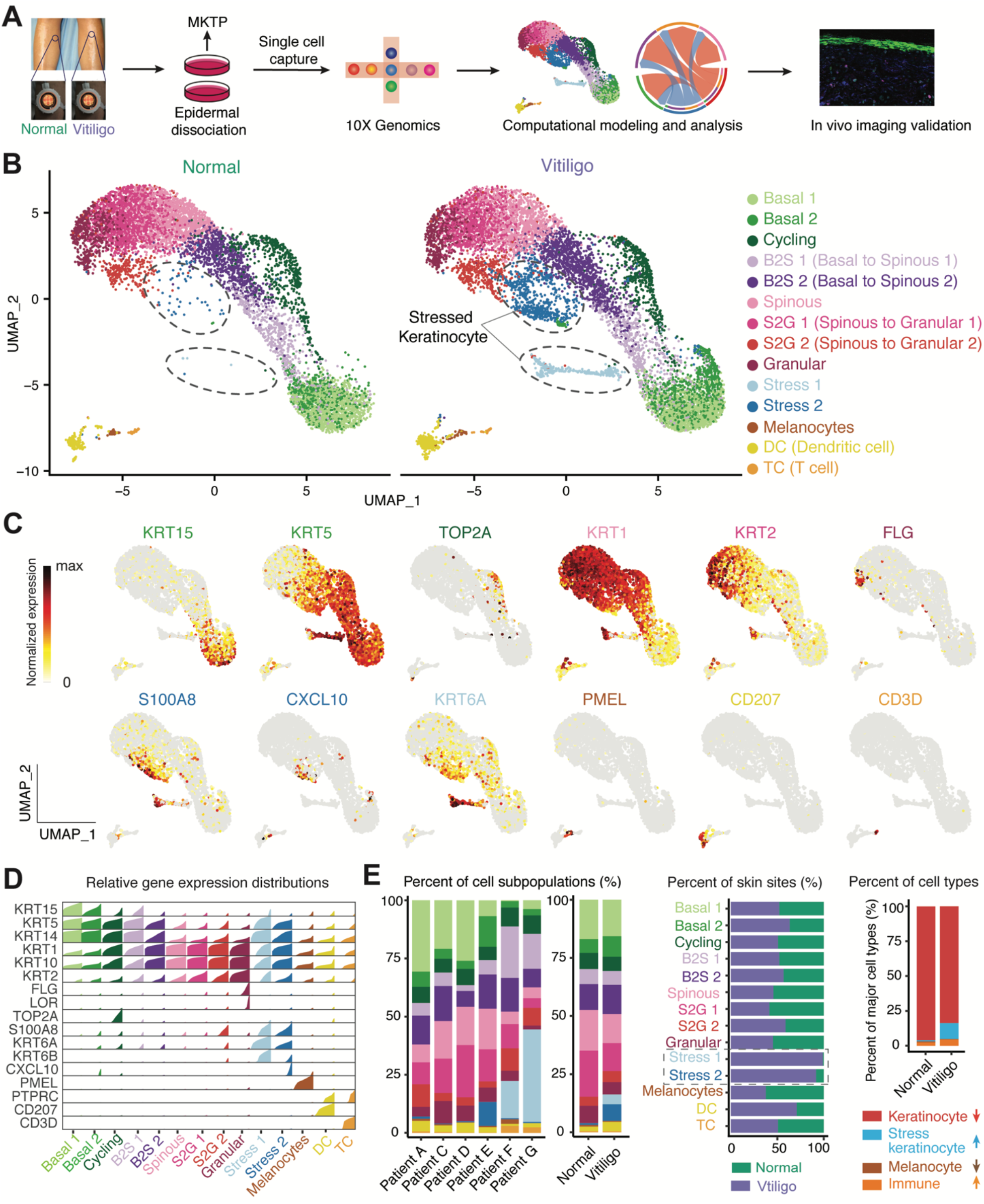
Single cell RNA-seq analyses of lesional and nonlesional skin reveal unique keratinocyte cell states in vitiligo patients. A) Schematic diagram of single cell isolation and scRNA-seq data analyses. B) UMAP plot of the cells from all patients in both nonlesional (left) and lesional skin (right). C) Feature plots showing expression of the selected markers in the UMAP space of all cells. D) High density plot showing relative gene expression of key marker genes in different cell subpopulations. Each density plot is composed by bar charts and bar height corresponds to the relative expression level of the gene in cells that is ordered from low to high. E) Percentages of cell subpopulations across patients, lesional and nonlesional skin (left). Comparison of the percentages of each cell subpopulation across lesional and nonlesional skin (middle). Comparison of the percentages of major cell types including keratinocytes, stress keratinocytes, melanocytes and immune cells across lesional and nonlesional skin (right). The bar plot shows that the percentages of keratinocytes and melanocytes decrease, while the percentages of stress keratinocytes and immune cells increase in lesional skin compared to nonlesional skin.

### Stress keratinocytes exhibit altered metabolism with dominant upregulation of OxPhos

To further characterize keratinocyte differences in detail between vitiligo lesional and nonlesional skin, we first performed differential expression analysis and found that lesional skin expressed higher levels of *KRT6A* and *KRT16* keratins that are not normally expressed in the mature interfollicular epidermis and are associated with insults like wounding and UV injury (fig. 3A) (*26*, *27*). Inflammatory and immune response related genes such as CD74, IFI27, IFI6 and IFITM1 were also significantly increased, which was further confirmed by the hallmark pathway enrichment analysis of the genes highly expressed in vitiligo lesional skin using the Molecular Signatures Database (MSigDB, fig. 3A) (*29*). In addition, we found that the top two enriched pathways were interferon gamma and alpha responses (fig. 3A), which is consistent with previous findings that lesional keratinocytes differed from their nonlesional counterparts in upregulation of interferon responses (fig. 3A) (*5*, *30*). Gene scoring analysis revealed downregulation of WNT signaling (fig. 3B, see Methods), consistent with the known role of WNT in melanocyte pigmentation (*5*, *30*). Since MPM demonstrated metabolic differences between nonlesional and lesional vitiligo skin, we further computed the signature scores of oxidative phosphorylation (OxPhos). Interestingly, higher scores were observed in lesional skin (fig. 3B).

**Fig 3.**
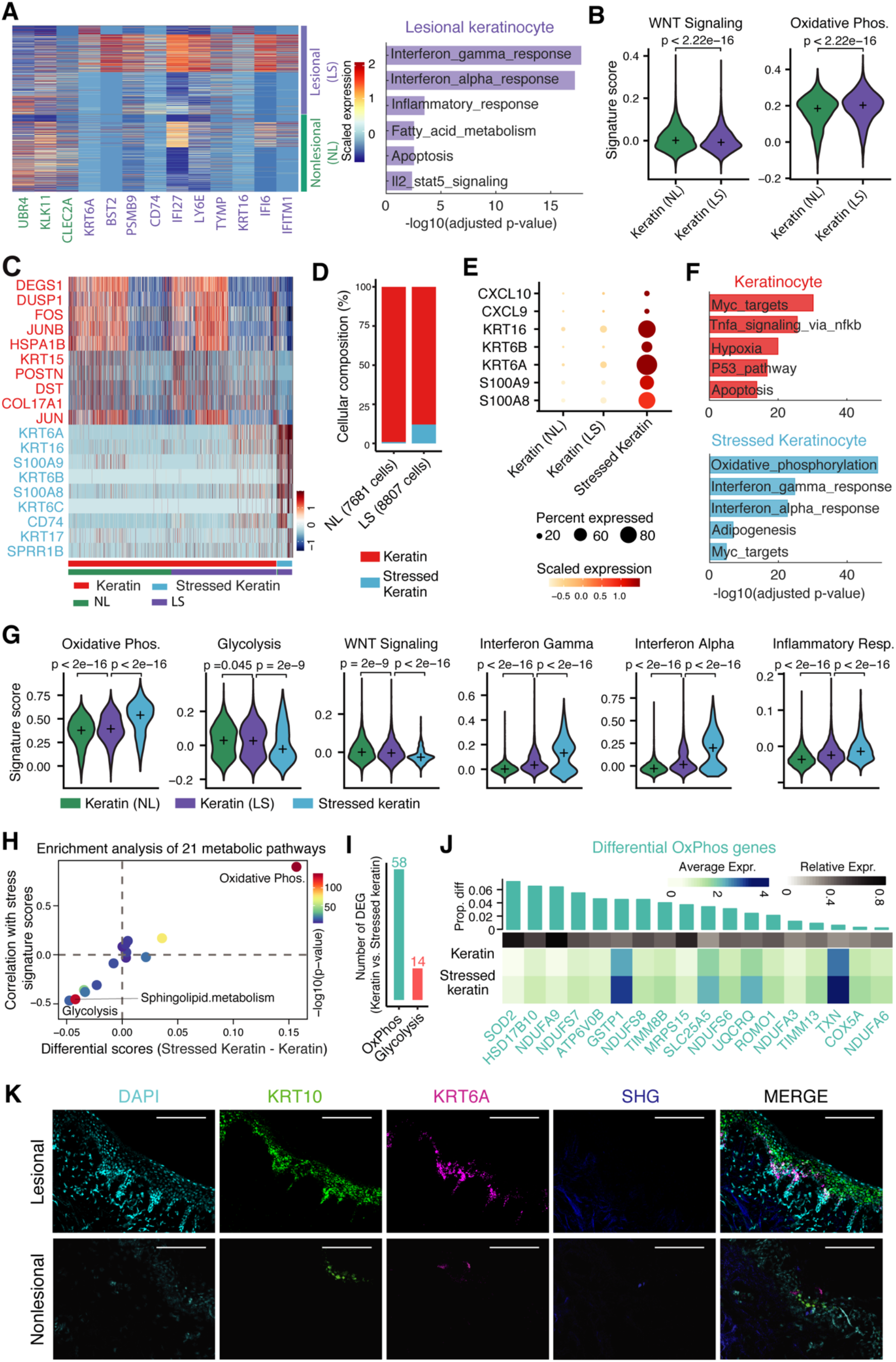
Stress keratinocytes have elevated metabolism and are the main source of CXCL9 and CXCL10. A) Heatmap of scaled expression levels of top 10 differentially expressed genes between nonlesional and lesional keratinocytes (left). Enriched Hallmark pathways of the highly expressed genes in lesional keratinocytes (right). B) Violin plots comparing signature scores of WNT signaling and OxPhos pathway between nonlesional and lesional skin. P-values are from two-sided Wilcox rank tests. C) Heatmap of scaled expression levels of differentially expressed genes between stressed keratinocytes and other keratinocytes. D) The composition of stressed keratinocytes and other keratinocytes in nonlesional and lesional skin. E) Dots plot of stress associated markers in nonlesional, lesional and stressed keratinocytes. The size represents the percentage of expressing cells and colors indicates the scaled expression. F) Enriched Hallmark pathways of highly expressed genes in stressed keratinocytes and other keratinocytes, respectively. G) Violin plots of signature scores of OxPhos, Glycolysis, WNT signaling, Interferon Gamma, Interferon Alpha and Inflammatory response across nonlesional, lesional and stressed keratinocytes. H) Enrichment analysis of 21 metabolic pathways in stress keratinocytes vs. other keratinocytes. Each dot represents one pathway. X-axis is the differential gene signature scores of each metabolic pathway between stressed keratin and other keratinocytes. Y-axis is the Pearson’s correlation of gene signature scores between each metabolic pathway and stress response. Gene signature scores of stress response were computed based on the marker genes of stressed keratinocytes. Colors represent the P-values from two-sided Wilcox rank tests of each gene signature score between stressed keratin and other keratinocytes. I) The number of differentially expressed OxPhos and Glycolysis-related genes in stress keratinocytes vs. other keratinocytes. J) Heatmap showing the average expression of top 18 differentially expressed OxPhos-related genes between stressed keratin and other keratinocytes. The top green bars represent the difference in the proportion of expressed cells between stressed keratin and other keratinocytes. K) RNAscope demonstrating Krt6A, Krt10 in situ hybridization in patient matched lesional and nonlesional punch grafting tissue. DAPI (cyan) was used to stain nuclei and second harmonic generation (blue) demonstrating location of collagen.

To figure out whether the above observed differences in signaling and metabolism were attributed to the unique stress keratinocytes in vitiligo lesional skin, we next focused on the difference between keratinocytes and stress keratinocytes. Differential expression analysis revealed distinct gene signatures between these two keratinocyte states (fig. 3C and F). In addition to *KRT6*, *KRT16*, *KRT17*, *S100A8* and *9* alarmins are known to be expressed in stress keratinocytes (fig. 3C) (*31*). Hallmark gene enrichment analysis of the differentially expressed genes showed that stress keratinocytes were enriched by OxPhos and interferon responses (fig. 3F). Since there were nearly no stress keratinocytes in nonlesional skin (fig. 3D), we focused on three keratinocyte groups: nonlesional keratinocytes, lesional keratinocytes and lesional stress keratinocytes. Comparison of these groups showed that CXCL9/10, KRT16, KRT6A/B and S100A8/9 were specifically expressed in stress keratinocytes instead of other two keratinocyte groups (fig. 3E). We further performed quantitative comparison of these three keratinocyte groups using gene scoring analysis (see Methods). Impressively, we observed dramatic differences between stress keratinocytes and both lesional and nonlesional keratinocytes, in terms of OxPhos, Glycolysis, WNT signaling, Interferon Gamma, Interferon Alpha and Inflammatory response (fig. 3F). Notably, significantly increased OxPhos and decreased glycolysis were consistent with our MPM imaging data (fig. 3F and fig. 1A). These results suggest that stress keratinocytes in vitiligo lesional skin dominantly account for the observed differences in signaling and metabolism between lesional and nonlesional skin.

To further examine whether OxPhos and glycolysis were the prominently impaired metabolic processes in vitiligo lesional skin, we quantitively evaluated the enrichment of 21 metabolic pathways using gene scoring analysis. We observed that OxPhos and Glycolysis were the most significantly altered pathways among all 21 metabolic pathways, which showed largest differences between stress keratinocytes and other keratinocytes and strongest correlations with stress signatures (fig. 3G). Of note, OxPhos and Glycolysis were highly positively and negatively correlated with stress signatures, respectively. There are 58 and 14 differently expressed OxPhos and Glycolysis related genes between stress keratinocytes and other keratinocytes (fig. 3H). Stress keratinocytes were enriched for genes associated with OxPhos, including SOD2, NDUFA9 and ATP6V0B. In contrast, keratinocytes expressed higher levels of genes associated with Glycolysis, including ALDH3A2, SDC1 and HSPA5. These results, combined with MPM data, indicate that a subpopulation of cells in vitiligo skin have altered metabolism and upregulate of OXPHOS.

We then performed RNAscope on patient-matched lesional and nonlesional skin to localize this keratinocyte population using KRT6A as it is highly expressed in this population (fig. 2C). We found that consistent with our MPM imaging, KRT6A expressing cells were enriched in the basal layer of the epidermis (fig. 3K).

### Analysis of cell-cell communication reveal major signaling changes in response to vitiligo

To systematically detect major signaling changes in stable vitiligo lesions, we applied our recently developed tool CellChat (*32*) to the scRNA-seq data of both nonlesional and lesional skin (see Methods). We observed almost twice number of interactions in lesional skin than nonlesional skin (fig. 4A). To study the prominent signaling pathways that contribute to the dysfunctional signaling in lesional skin, we compared each signaling pathway between nonlesional and lesional skin using the concept of information flow defined as a sum of the communication probability among all pairs of cell groups. We found that several pathways were only activated in nonlesional skin (fig. 4B), including WNT, PTN and VEGF, consistent with the role of WNT activation in regulating melanocyte differentiation(*33*). In contrast, many inflammatory pathways prominently increase their information flow at lesional skin as compared to nonlesional skin, such as CXCL, IL4, IL6, LT, LIGHT, TWEAK, TNF, VISFATIN and GALECTIN.

**Fig 4.**
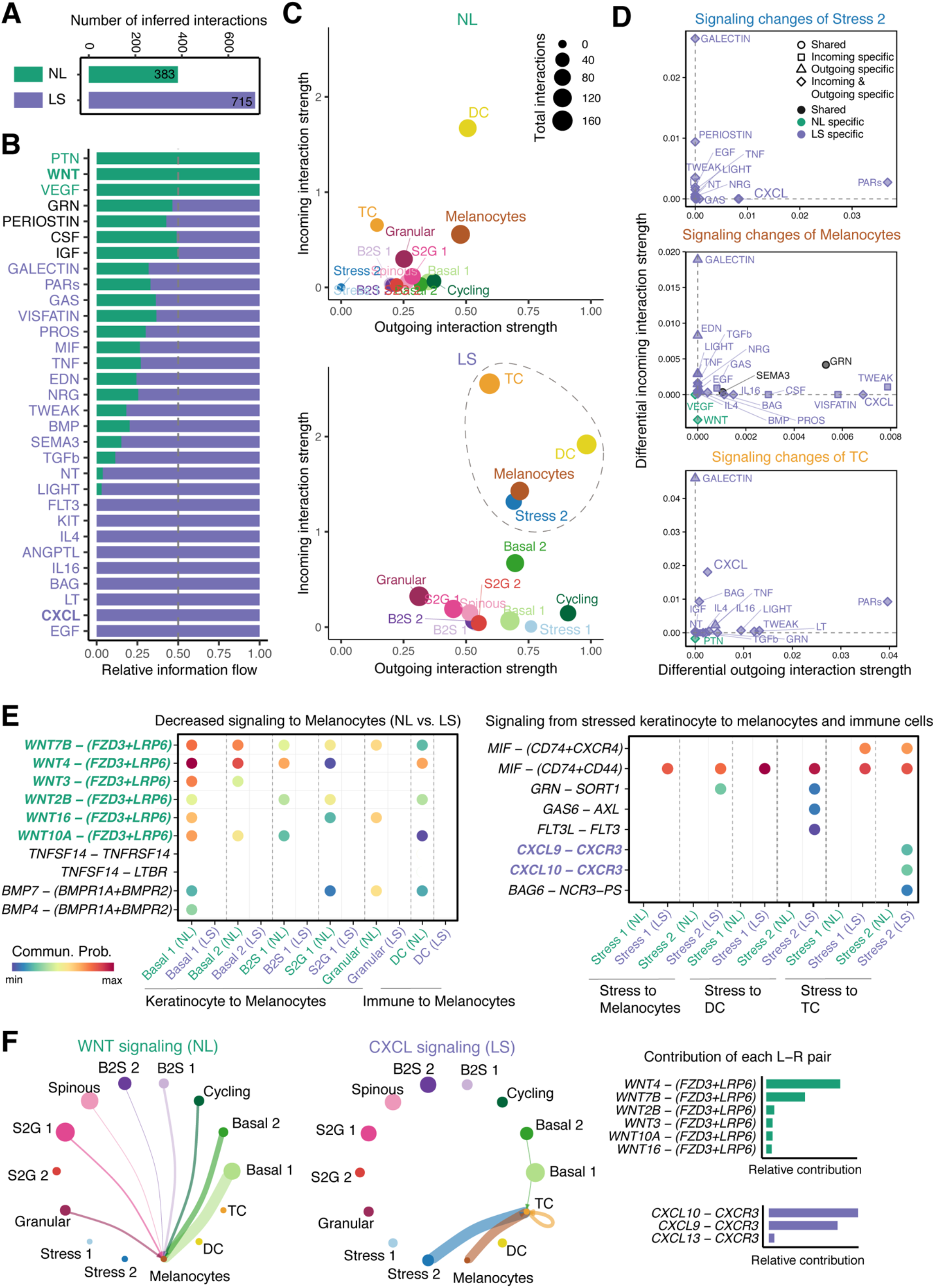
Cell-cell communication analysis reveals major signaling changes between nonlesional and lesional vitiligo skin. A) Number of inferred interactions among all cell subpopulations between nonlesional (NL) and lesional (LS) skin. (B). The relative information flow of all significant signaling pathways within the inferred networks between nonlesional and lesional skin. Signaling pathways labeled in green represent pathways enriched in nonlesional skin, the middle ones colored by black are equally enriched in both nonlesional and lesional skin, and the ones colored by purple are more enriched in lesional skin. (C) Visualization of outgoing and incoming interaction strength of each cell subpopulation in the inferred cell-cell communication network of nonlesional (top) and lesional skin (bottom). The dot sizes are proportional to the number of total interactions associated with each cell subpopulation. Dashed circle indicates the most altered cell subpopulations when comparing the outgoing and incoming interaction strength between nonlesional and lesional skin. (D) The signaling changes associated with the three most altered cell subpopulations. (E) Bubble plot in left panel shows the decreased signaling from keratinocyte and immune subpopulations to melanocytes (nonlesional vs. lesional skin). Bubble plot in right panel shows all significant signaling from stress keratinocyte to melanocytes and immune subpopulations. (F) Inferred cell-cell communication networks of WNT and CXCL signaling in nonlesional and lesional skin, respectively (left). The edge width is proportional to the inferred communication probabilities. The relative contribution of each ligand-receptor pair to the overall signaling pathways (right).

We next studied how different cell subpopulations changed their signaling patterns from nonlesional to lesional skin using network centrality analysis, which computes the outgoing and incoming interaction strength of each subpopulation to represent the likelihood as signaling sources and targets, respectively. This analysis revealed that T cells emerged as major signaling targets while dendritic cells (DC) became dominant signaling sources. Melanocytes and Stress 2 keratinocytes also prominently increased their outgoing and incoming signaling from nonlesional to lesional skin (fig. 4C). We then asked which signaling pathways contributed to the signaling changes of these populations. Differential interaction analysis showed that the prominently increased outgoing signaling of Stress 2 keratinocytes and Melanocytes and the incoming signaling to T cells was CXCL (fig. 4D), suggesting that CXCL signaling pathway was the dominantly dysfunctional signaling sent from Stress 2 keratinocytes and Melanocytes to T cells. Of note, WNT is the major decreased incoming signaling of Melanocytes.

By studying the signals sent to melanocytes, we found that a relative deficiency of WNT and BMP signaling was noted in keratinocytes and DC in lesional skin. In particular, WNT signal was seen in all keratinocyte populations in nonlesional skin with WNT4 and WNT7B driving the signaling (fig. 4E, F). For the signaling from stress keratinocyte to melanocytes, DC and TC cells, MIF and CXCL signaling were highly active in lesional skin. Notably, for the signaling from stress keratinocyte to TC, ligands CXCL9 and CXCL10 and their receptor CXCR3 were found to be uniquely active in lesional skin (fig. 4E, F). Taken together, our analyses indicated the prominent alteration of cell-cell communication networks in vitiligo lesional skin and predicted major signaling changes that might drive vitiligo pathogenesis.

### Pseudotemporal dynamics reveal transition dynamics of stress keratinocytes

To explore the role of stress keratinocytes in keratinocyte differentiation, we performed pseudotemporal trajectory analysis using all keratinocyte cells except for cycling cells from all samples. By applying the diffusion-based manifold learning method PHATE (*34*, *35*) to the batch-corrected data obtained from scMC(*25*), we observed a differentiation path in the nonlesional skin, recapitulating sequential stages of keratinocyte differentiation process from basal state to terminally differentiated granular state. However, in vitiligo lesional skin, in additional to the known keratinocyte differentiation path (Path 1), another potential differentiation path (Path 2) was found to attribute to stress keratinocytes (fig. 5 A). Using an unsupervised pseudotemporal trajectory inference tool Monocle 3 (*36*), we showed the stress keratinocytes indeed contributed to alternative differentiation paths, indicating a transition from an early intermediate keratinocyte state (basal to spinous transition) to stress keratinocytes, to a late intermediate keratinocyte state (spinous to granular transition), and then to granular state (fig. 5 B). Such observation was further confirmed using another trajectory inference approach PAGA (*37*), showing strong likelihood of the transition between stress keratinocytes and the late keratinocyte states (fig. S6A). To further analyze the keratinocyte differentiation dynamics, we performed RNA velocity analysis using scVelo, a computational tool that can predict potential directionality and speed of cell state transitions based on levels of spliced and unspliced mRNA(*38*). RNA velocity analysis also provided evidence for enhanced transition dynamics from stress keratinocytes to the late keratinocyte state (fig. S6B). Together, in addition to the normal keratinocyte differentiation trajectory, these analyses showed the transition dynamics of stress keratinocytes contribute to an altered keratinocyte differentiation trajectory in vitiligo lesional skin.

**Fig 5.**
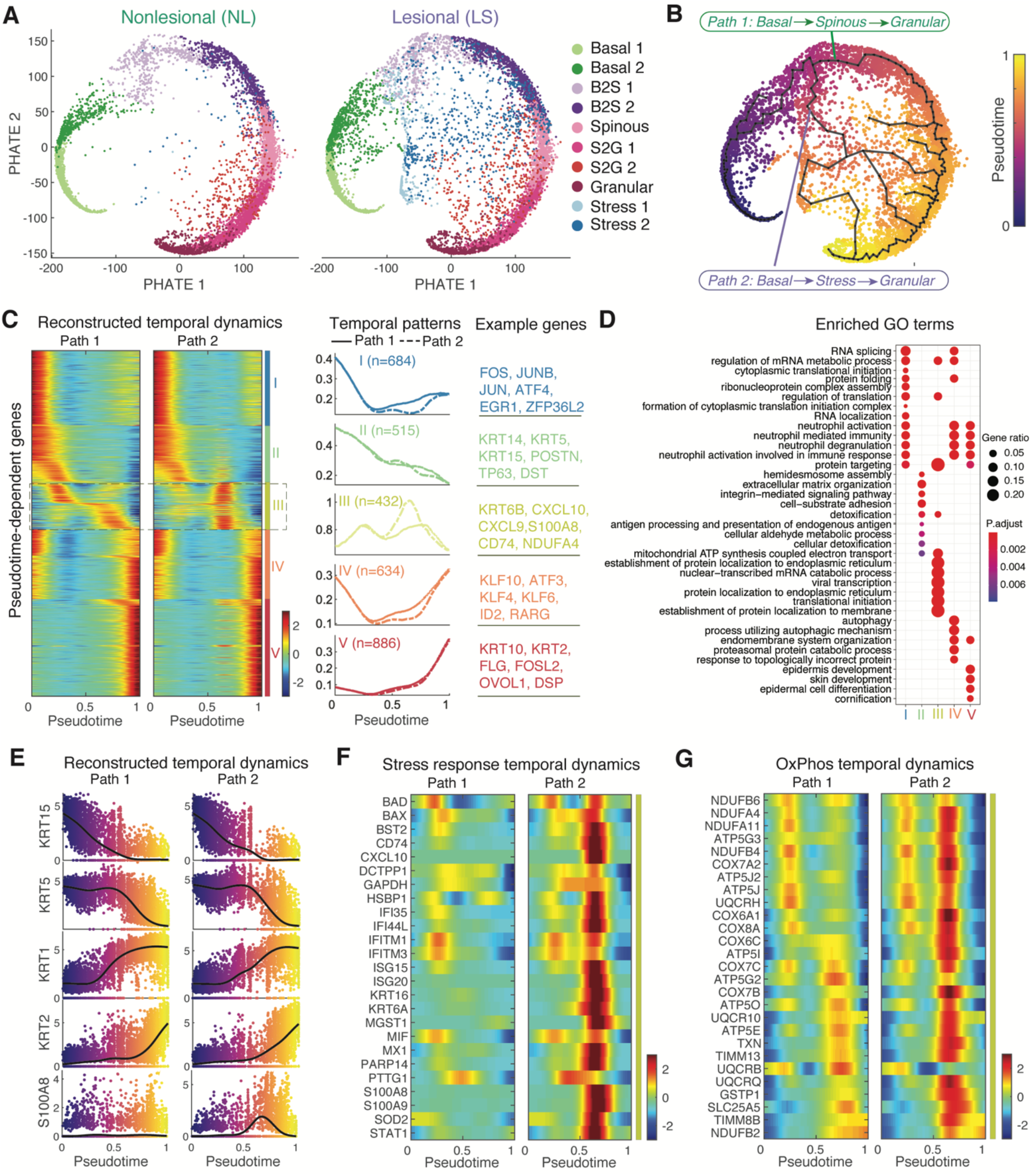
Pseudotemporal dynamics reveal transition dynamics of stress keratinocytes. A) Projection of keratinocytes onto the PHATE space revealed the potential lineage relationships between different keratinocyte subpopulations in nonlesional (NL, left panel) and lesional (LS, right panel) skin. Cells were colored by the annotated cell identity. (B) The inferred pseudotemporal trajectories of all cells using Monocle 3. Cells were colored by the inferred pseudotime. Pseudotemporal trajectory analysis revealed two potential transitional paths, as indicated by Path 1 and Path 2. (C) Pseudotemporal dynamics of all pseudotime-dependent genes along the Path 1 and Path 2. Each row (i.e., gene) is normalized to its peak value along the pseudotime. These genes were clustered into five groups with the average expression patterns (middle) and representative genes (right). Solid and dashed lines indicate the average expression of a particular gene group in Path 1 and Path 2, respectively. The number of genes in each gene group is indicated in parenthesis. (D) Enriched biological processes of the five gene groups in (C). (E) The reconstructed pseudotemporal dynamics of selected marker genes along the inferred pseudotime in Path 1 and Path 2, respectively. Black lines represent the average temporal patterns that were obtained by fitting a cubic spline. Cells were colored by the inferred pseudotime. (F-G) Pseudotemporal dynamics of the pseudotime-dependent genes related with the stress response and OxPhos along the inferred pseudotime in Path 1 and Path 2, respectively.

We next sought to identify key molecular changes that may be important for keratinocyte cell state transitions using scEpath (*35*). scEpath identified 1284 and 3151 pseudotime-dependent genes over the normal (Path 1) and alternative keratinocyte differentiation trajectories (Path 2), respectively (fig. 5C). These pseudotime-dependent genes were further classified into five groups based on their pseudotemporal dynamics. Interestingly, the gene group III exhibited distinct expression dynamics along the Path 1 versus Path 2 while the remaining gene groups followed very similar dynamical trends on both trajectories. Genes in Group III included not only stress keratinocyte-related signatures such as KRT6B, CXL10, CXCL9, S100A8 and CD74, but also OxPhos-associated signatures such as NDUFA4 and ATP5G3 (fig. 5C). Further GO enrichment analysis revealed distinct enriched biological processes among these five gene groups, including the enriched metabolic processes in group III (fig. 5D). The reconstructed pseudotemporal dynamics of typical maker genes well recapitulated the expected keratinocyte differentiation dynamics (fig 5E). As expected, we observed stronger activation of stress response, inflammatory response and OxPhos associated genes in the Path 2 compared to Path 1 (fig. 5F-G). Taken together, stress keratinocytes induce an altered keratinocyte differentiation trajectory with strong activation of inflammatory response and OxPhos related gene expression in vitiligo lesional skin.

## DISCUSSION

To date, the study of human vitiligo and cell-cell interactions in the tissue microenvironment (TME) have largely been limited to traditional *in vitro* cultures and immunohistochemistry methods due to the lack of tools to assess cellular changes *in situ*. Here, we combine MPM *in vivo* imaging of stable vitiligo patients and various scRNA-seq analyses to demonstrate that a small subpopulation of stress keratinocytes in the basal/parabasal layer exhibit a unique signature – metabolic preferences for oxidative phosphorylation, expression of stress keratins, alarmins and CXCL9/10 and diminished WNT signaling – and likely drive the persistence of white patches in vitiligo. Our data suggest that it is feasible to use MPM as a noninvasive method to track metabolically altered populations of keratinocytes in vitiligo. Previous studies on metabolic alterations in vitiligo largely focused on melanocytes’ increased susceptibility to oxidative insults such as H_2_O_2_ due to decreased expression of antioxidant pathways (*39*–*41*) Oxidative stress led to HMGB1 release by cultured melanocytes, which then stimulates cytokine release by keratinocytes(*42*). Studies on cultured keratinocytes from vitiligo skin showed swollen mitochondria and similar increased susceptibility to oxidative stress (*11*, *43*). However, definitive studies looking at keratinocyte metabolism and its contributions to vitiligo have been lacking. Our study addresses this gap and demonstrates that specific basal and parabasal keratinocyte states exhibit increased OXPHOS and communicate with T cells via the CXCL9/10/CXCR3 axis and exhibit decreased WNT signaling to melanocytes.

Most studies on vitiligo have focused on active disease and the importance of the CXCL9/10/CXCR3 axis is well established from studies on human skin samples (*4*, *8*, *30*, *44*). Stable vitiligo, however, remains enigmatic(*45*). Transcription analyses on depigmented whole skin shows minimal immune activation with no *CXCL10* elevation (*5*). Flow cytometry of stable vitiligo skin blisters demonstrated the presence of a small population of melanocyte-specific CD8^+^ resident memory T cells (T_RM_) and depletion of T_RM_ by targeting CD122 led to re-pigmentation in a mouse model of vitiligo (*46*). By using scRNA-seq to identify changes in cellular compositions in stable vitiligo skin, we identified a keratinocyte state with transcriptome changes important in communicating with other cell types to drive disease persistence. The signals from stress keratinocytes were likely lost from averaging cell gene expression in previous whole skin transcriptional studies, accounting for observed differences in CXCL10 expression in our study (*5*, *47*, *48*). By utilizing CellChat analyses, our data highlights that in stable vitiligo, a small epidermal niche of metabolically altered stress keratinocytes communicate with T cells and melanocytes to form local inflammatory circuits to drive disease persistence (Fig. 6), highlighting vitiligo involves multiple etiologic factors(*49*). Keratinocytes as drivers of local inflammatory loops have been suggested in atopic dermatitis and psoriasis (*40*). We show that similar loops are important in vitiligo persistence but how stress keratinocytes are established in the first place and whether they play a key role in the maintenance of this cellular circuitry remain obscure. Are stress keratinocytes a consequence of intrinsic keratinocyte differentiation defects in genetically susceptible individuals? Or do extrinsic signals from other tissue cells (T cells, the absence of melanocytes or a combination) drive this cellular state? Studies are currently underway to investigate when metabolically altered keratinocytes first appear and how they may affect the repigmentation process in patients undergoing treatment.

**Figure 6.**
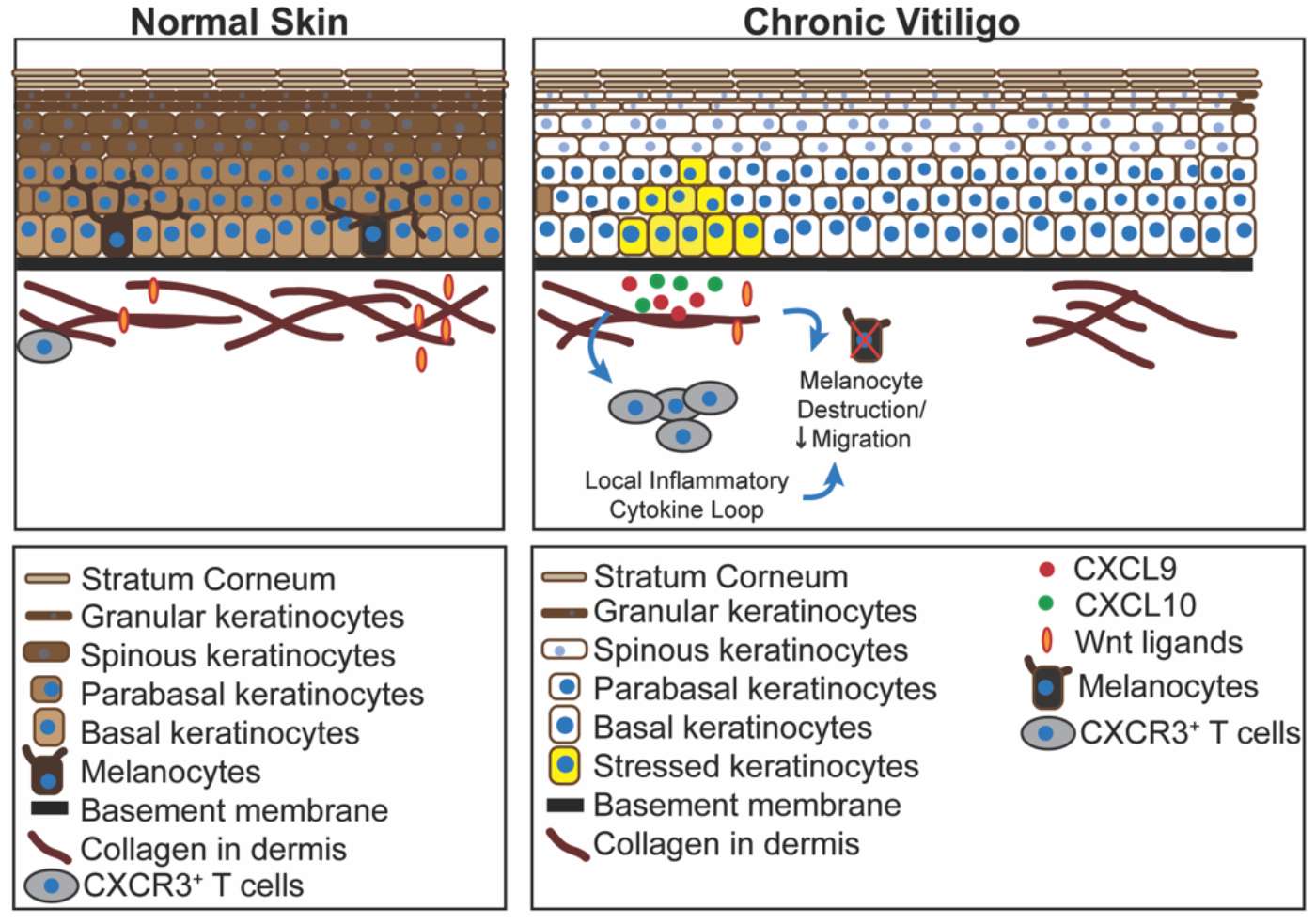
Model of stable vitiligo as an active disease state. Stress keratinocytes, which exhibit a preference for oxidative phosphorylation, are enriched in vitiligo skin and secrete CXCL9 and 10 to communicate with T cells to form local inflammatory circuits. They express lower levels of Wnt ligands, which have been shown to play a role in melanocyte migration.

The findings of our study raise the possibility of targeting keratinocyte metabolism in vitiligo treatment. Intriguingly, biguanides such as phenformin and metformin that inhibit oxidative phosphorylation have been shown to affect keratinocyte differentiation and pigmentation (*50*, *51*). Whether these drugs will also inhibit keratinocyte-derived signals that affect immune cell and melanocyte recruitment is unclear and represent an unexplored area for drug targeting in vitiligo. Interestingly, stress keratinocytes expressing *KRT6*, *16* and *S100A8/9* have been identified in the human epidermis of psoriasis and melanomas, raising the possibility that they can play a wide variety of roles in the diseased skin tissue microenvironment (*52*–*54*). Further studies on stress keratinocytes will improve our understanding how keratinocyte states affect the tissue microenvironment and contribute to disease pathogenesis.

A caveat to our study is that scRNA-seq analyses were performed on skin blisters which do include fibroblasts and other dermal cell types. We chose blisters as they represented a nonscarring method to collect vitiligo skin samples and had previously been shown to be sufficient to predict disease activity (*4*). The absence of dermal tissue in our analysis may account for the lack of innate immune cells that other groups have identified(*55*, *56*). Detailed analysis of vitiligo skin has been hampered by the lack of fresh tissue samples for analysis, as induction of blisters or biopsy itself can induce the disease (*1*). Moreover, patient to patient variation in vitiligo can be significant, which makes it difficult to make generalized conclusions on the pathogenesis of the disease. Here we have coupled together imaging of lesional and non lesional skin with single cell sequence analysis that specifically controls for sample to sample and patient to patient variability (scMC) to make generalizable conclusions regarding disease pathogenesis, providing a roadmap for the study of other diseases that are controlled by cell-cell interactions in tissue.

Our data indicate that stress keratinocytes have altered metabolic preferences, drive local inflammation in the skin microenvironment and can be visualized *in situ* in human patients using noninvasive MPM imaging. These results are significant because they provide evidence for a potential link between stress keratinocytes and vitiligo persistence. They also indicate that MPM imaging can also be used to follow vitiligo patients longitudinally to better understand the role stress keratinocytes in disease pathogenesis and identify areas that could be targeted by new therapies. These new therapies could range from targeted destruction of altered keratinocytes (laser therapies) or pharmacologic modulation of their physiology. As an example, our work implicates the combination of therapies that reverse keratinocyte metabolic defects and JAK inhibitors as a novel treatment for vitiligo. Studying this process will require the generation of new tissue models to study vitiligo pathogenesis that can overcome the limitations of mouse models, where tripartite interactions between melanocytes keratinocytes and immune cells in the epidermis is largely absent.

## MATERIALS AND METHODS

### Study Design

This study utilized noninvasive MPM and scRNA-seq to study patient-matched lesional vs. nonlesional skin in stable vitiligo and how intercellular communications are affected in depigmented skin. Imaging, suction blister and punch skin biopsy of patients were performed under IRB-approved protocols at UC Irvine and samples were de-identified before use in experiments. Vitiligo skin samples were obtained after examination by board-certified dermatologists (JS, AKG). Control skin was acquired from tumor excision tips without notable pathology from patients without vitiligo. Stable vitiligo lesions were characterized by the absence of koebnerization, confetti-like depigmentation or trichome lesions and those that have not grown in size for at least one year (*24*). Non-lesional sites were selected as normal-appearing, non-depigmented skin on the thigh when examined by Wood’s lamp.

### Patients for Imaging

Twelve vitiligo patients and five volunteers with normal skin were imaged *in vivo* by MPM. All vitiligo patients had stable vitiligo, defined by no change in size for at least one year and do not exhibit features of active vitiligo such as koebnerization, confetti-like depigmentation and trichome (*24*). Patients were previously unresponsive to past treatment attempts, and had no treatment in the three months before imaging for this study. Vitiligo patient ages were 34-74 with an average age of 56. Vitiligo lesion locations included wrist (2), hand (2), leg (5), arm (1), face (1), and neck (1). Nonlesional pigmented skin was selected after Wood’s lamp exam on separate body sites or at least 12cm from closest depigmented macule. All *in vivo* measurements were conducted according to an approved institutional review board protocol of the University of California, Irvine (HS No. 2018-4362), with written informed consent obtained from all patients.

### MPM imaging

We used an MPM-based clinical tomograph (MPTflex, JenLab, GmbH, Germany) for the in vivo imaging of the vitiligo and normal skin. This imaging system consists of a femtosecond laser (Mai Tai Ti:Sapphire oscillator, sub-100 fs, 80 MHz, tunable 690– 1020 nm; Spectra-Physics), an articulated arm with near-infrared optics, and beam scanning module. The imaging head includes two photomultiplier tube detectors used for parallel acquisition of two-photon excited fluorescence (TPEF) and second harmonic generation (SHG) signals. The excitation wavelength used in this study was 760 nm. The TPEF and SHG signals were detected over the spectral ranges of 410 to 650 nm and of 385 to 405 nm, respectively. We used a Zeiss objective (40×, 1.3 numerical aperture, oil immersion) for focusing the laser light into the tissue. The laser power used was 5 mW at the surface and up to 30 mW in the superficial dermis of the skin. We acquired the MPM data as z-stacks of en-face images from the stratum corneum to the superficial dermis. The field of view (FOV) for each optical section was 100 × 100 μm^2^ and the step between the optical sections was 5 μm. We imaged the patients’ vitiligo lesional area, and a normally pigmented area on the upper thigh as control. The rationale for selecting the thigh location as control site for imaging was based on to the fact that the patients we imaged, being unresponsive to prior treatment of vitiligo, were scheduled for micrografting therapy. Imaging locations for healthy volunteers with normal skin were the sun exposed dorsal forearm, and the non-sun exposed volar upper arm to focus on areas with relatively higher pigment amounts (sun-exposed), and relatively lower pigment amounts (non sun-exposed). Due to the limited FOV of each individual scan, we acquired several stacks of images within each site in order to sample a larger area. Thus, a total of 1,872 images were acquired for this study, corresponding to an average of 18 images for each imaging site. Images were 512 × 512 pixels and were acquired at approximately 6 s per frame. All images were color-coded such that green and blue represent the TPEF and SHG signals, respectively. In MPM imaging of skin, the contrast mechanism is based on two-photon excited fluorescence (TPEF) signal from NADH, FAD, keratin, melanin, and elastin fibers (*57*–*59*) and on second harmonic generation (SHG) signal from collagen(*60*). These images were used as a basis for the mitochondrial clustering analysis (see supplementary methods).

### Suction Blister Induction and cell isolation for single-cell RNA sequencing

All procedures were conducted according to an approved institutional review board protocol of the University of California, Irvine (HS No. 2018-4362), with written informed consent obtained from all patients. The donor skin sites were cleaned with ethanol wipes and 5 suction blisters (1cm diameter) were created by applying a standard suction blister device. We unroofed the blisters and used half for melanocyte-keratinocyte transplant procedure(*61*). The rest of the blisters were incubated in trypsin for 15 minutes at 37°C, followed by mechanical separation and centrifugation at 1000 rpm for 10 minutes at 4°C to pellet cells. Cells were washed with 0.04% UltraPure BSA:PBS buffer, gently re-suspended in the same buffer, and filtered through a 70μm mesh strainer to create a single cell suspension. Cells were washed and viability was calculated using trypan blue. scRNA-seq was performed by the Genomics High Throughput Sequencing facility at the University of California, Irvine with the 10x Chromium Single Cell 3’ v2 kit (10x Genomics). None of the patients that were imaged overlapped with the cohort of patients that were analyzed by single cell RNA sequencing. Details of the single cell data analysis is provided in the supplementary methods.

### Patient Samples for RNAscope

All procedures were conducted according to an approved institutional review board protocol of the University of California, Irvine (HS No 2018-4362) with written informed consent obtained from all patients. Briefly, 2mm biopsies were performed on lesional and nonlesional skin as part of punch grafting treatment for three patients. Control skin was acquired from tumor excision tips without notable pathology from patients without vitiligo. Skin samples were immediately frozen and embedded in OCT. Tissues were stored at −80°C and cryosections (10mm thick) of skin were collected on Fisherbrand Superfrost Plus microscope slides. Sections were dried for 60-120 minutes at −20°C then used immediately or within 10 days. *In situ* hybridization was performed according to the RNAscope Multiplex Fluorescent Reagent Kit v2 (Cat. No. 320293).

Briefly, slides were fixed in cold 4% PFA for 15 minutes then dehydrated in 50%, 70%, and 100% ethanol for 5 minutes each at room temperature (RT). H_2_O_2_ was applied for 10 minutes at RT and treated with protease IV for 30 minutes. C2 and C3 probes were diluted in C1 probes at a 1:50 ratio and incubated for 2 hours at 40°C. C1 probes were detected with TSA-fluorescein (Akoya Biosciences), C2 probes with Opal-620 and C3 probes with Opal-690 (Akoya Biosciences). Before mounting, DAPI was added to label the nuclei. Images were acquired using a Leica SP8 FALCON/DIVE (20x objective, 0.75 NA).

### Statistical Analysis

Statistical comparisons of median β and β variability were conducted using linear mixed effects models in SAS JMP Pro 14. Variables such as patient number and imaging location were modeled as random effects. Whether an area of skin was lesional or non-lesional was modeled as a fixed effect when comparing metrics of mitochondrial clustering among patients. Whether an area of skin was sun-exposed or non-sun-exposed was modeled as a fixed effect when comparing metrics of mitochondrial clustering among healthy volunteers. The significance level for all statistics was set to α = 0.05.

## Supplementary Materials

Supplementary Materials and Methods

Fig. S1. Mitochondrial Clustering Distributions for Basal Keratinocytes.

Fig. S2. Quality control metrics of the data.

Fig. S3. The Difference between DC and TC.

Fig. S4. Analysis results of scRNA-seq data of normal skin.

Fig. S5. Analysis results of scRNA-seq data of all patients by Seurat.

Fig. S6. Pseudotime analysis results of scRNA-seq data of all patients.

Table S1. Clinical Characteristics of Stable Vitiligo Patients for MPM and RNAscope

Table S2. Clinical Characteristics of Stable Vitiligo Patients for scRNA-seq

## Acknowledgments

This work was made possible, in part, through access to the Genomics High Throughput Facility Shared Resource of the Cancer Center Support Grant (P30CA-062203) at the University of California, Irvine and NIH shared instrumentation grants 1S10RR025496-01, 1S10OD010794-01, and 1S10OD021718-01. The content is solely the responsibility of the authors and does not necessarily represent the official views of the National Institutes of Health.

## Funding

NIH grant 5KL2TR1416-6 (JS)

NIH grant R21AR073408 (AKG)

NIH grant U01AR073159 (QN)

NSF grant DMS1763272 (QN)

Simons Foundation grant (594598, QN)

NIH/NIAMS – P30-AR075047 (LZ, JLF, SJ)

NIH/NCI – U54-CA217378 (LZ, JLF, SJ)

## Author contributions

Conceptualization: MB, QN, AKG, IG, BJT
Methodology: JS, GL, LZ, SJ, JLF, CM, CP, FRD, PM
Investigation: JS, GL, LZ, SJ, JLF, SJJ, CP, CM, JK, FRD, PM
Visualization: JS, GL, LZ, SJ, JLF, CP
Funding acquisition: JS, MB, QN, IG, AKG
Project administration:
Supervision: IG, MB, QN, AKG, BJT
Writing – original draft: JS, GL, LZ
Writing – review & editing: JS, GL, LZ, SJ, JLF, SJJ, CP, IG, QN, MB, AKG

## Competing interests

Authors declare that they have no competing interests

## Code availability

Code for the scRNA-seq analysis have been deposited at the GitHub repository (https://github.com/amsszlh/Codes_for_paper_scRNA-seq_vitiligo)

scMC is publicly available as an R package under the GPL-3 license. Source codes, tutorials, and reproducible benchmarking codes have been deposited at the GitHub repository (https://github.com/amsszlh/scMC) and Zenodo repository (DOI: https://doi.org/10.5281/zenodo.4138819).

CellChat is publicly available as an R package. Source codes, as well as tutorials have been deposited at the GitHub repository (https://github.com/sqjin/CellChat). The web-based CellChat Explorer, including Ligand-Receptor Interaction Explorer for exploring the ligand-receptor interaction database and Cell–Cell Communication Atlas Explorer for exploring the intercellular communications in tissues, is available at http://www.cellchat.org/.

## Data and materials availability

Data has been submitted the GEO data base, awaiting accession number.

## Notes

### Competing Interest Statement

The authors have declared no competing interest.

