## Supplemental materials for "Multimodal Analyses of Stable Vitiligo Skin Identify Tissue Interactions That Control Disease Persistence"

### Supplementary Materials

### Supplementary Materials and Methods

#### Mitochondrial Clustering

All image processing steps were performed in MATLAB using an approach established previously(*21*). Several masks were created and combined in order to isolate cytoplasmic autofluorescence. An SHG mask was primarily created to remove contributions from collagen and stromal autofluorescence at the interface of the epidermis and dermis. Contrast-limited adaptive histogram equalization (CLAHE) was applied to SHG images and features were subsequently segmented using Otsu's global thresholding. The SHG mask was finalized by applying a median filter to remove noise and taking the complement of the image to mask features corresponding to the segmented signal. Features corresponding to highly autofluorescent biomolecules such as keratin and melanin were masked using similar methods. CLAHE was applied to TPEF images and an Otsu's global threshold was calculated. Pixels with intensity values 1.5X greater than the Otsu's global threshold were segmented and masked. This empirically determined threshold was applied to all optical sections and was determined based on the propensity to remove highly autofluorescent signatures without masking pixels from intermediate cell layers which would not contain fluorophores such as keratin and melanin. The removal of nuclear and interstitial regions was achieved by applying 3 serial bandpass filters to contrast-limited adaptive histogram equalized TPEF images. Remaining features were segmented using Otsu's global thresholding. A circular mask with a 500-pixel diameter was created to remove dim image corner artifacts. Masks were finalized with the removal of objects less than 8-pixels in size. The final mask was applied to raw TPEF images, which were then subjected to a digital object cloning (DOC) process. The DOC process randomly fills any void pixels from the masking process with signal that was identified as cytoplasm. No pixels are overwritten during this process and it is replicated 5 times. The average power spectral density (PSD) of the 5 DOC images was then computed and fitted with an equation of the form $R(k) = Ak^{-\beta}$ for spatial frequencies (k) less than 0.118 $\mu m^{-1}$ (features smaller than 8.5 μm). The absolute value of the fitted exponent, β, represents the degree of mitochondrial clustering within the cytoplasm. Mitochondrial clustering was computed for optical sections ranging from the stratum corneum to the stratum basale. Depth-dependent metrics of mitochondrial clustering were computed for each stack of images. β variability represents the sample variance of β values as a function of depth and aims to capture depth-dependent changes in metabolism. Median β represents the median β value as a function of depth and aims to capture the overall level of metabolic activity.

Mitochondrial clustering was calculated in the same manner for single-cell analysis. Due to the relatively low levels of contrast in the basal layer of the epidermis, single cells had to be manually segmented. One optical section per patient region of interest was segmented for single cells. Approximately 5 – 10 single cells were masked per image. A total of 182 cells from lesional and 258 cells from non-lesional regions were included for analysis. All vitiligo patients included in the imaging studies were represented in the total cell populations. The heterogeneity level of the corresponding distributions was quantified using a previously established heterogeneity index(*60*), based on fitting a 2-Gaussian mixture model to each distribution. Briefly, the heterogeneity index,

H, can be computed using the equation $H = -\sum d_i p_i \ln p_i$, where $i$ denotes each subpopulation, $d$ denotes the absolute value of the difference between the median of a subpopulation and the median of the total population, and $p$ denotes the Gaussian mixing proportion of the subpopulation. 2-Gaussian mixture models were derived using SAS JMP Pro 14 statistical software.

**Processing and quality control of scRNA-seq data**

Sequencing libraries were prepared using the Chromium Single Cell 3/ v2 protocol (10x genomics). Sequencing was performed on Illumina HiSeq4000 platform (Illumina). FASTQ files were aligned utilizing 10x Genomics Cell Ranger 2.1.0. Each library was aligned to an indexed hg38 genome using Cell Ranger Count. Cell Ranger Aggr function was used to normalize the number of mapped reads per cells across the libraries. Patient B sample and lesional skin of Patient G sample did not have enough viable cells and was excluded from further analysis (table S1). Quality control parameters were used to filter cells with 200-4000 genes with a mitochondrial percentage under 18% for subsequent analysis.

**Integation and clustering analyses of scRNA-seq data**

Integration and clustering of cells was performed using the scMC R package, which is a R toolkit for integrating and comparing multiple scRNA-seq experiments across different conditions. And scMC learns a corrected matrix, which is a shared reduced dimensional embedding of cells that preserves the biological variation while removing the technical variation(*25*). The data of each lesional and nonlesional skin of each patient were treated as one condition. Therefore, the input of the scMC is a list with 11 elements, with each element being one condition. The parameters used for this data are shown as follows: *resolution =1*; *quantile.cutoff = 0.5*; *similarity.cutoff = 0.65*. To identify cell clusters, principle component analysis (PCA) was first performed on the corrected matrix of scMC and the top 40 PCs with a resolution = 1 were used to obtaining 14 clusters for all the samples.

**Calculation of signature score of a gene set**

For gene scoring analysis, most gene sets were acquired from the MSigDB database (https://www.gsea-msigdb.org/gsea/msigdb/). Gene sets of metabolic pathways were from published literature (*61*). Specific genes in each gene set and their sources are listed in Table S3. The AddModuleScore function in Seurat R package was then used to calculate the signature score of each gene set in each cell. The two-sided Wilcoxon rank sum test was used to evaluate whether there are significant differences in the computed signature scores between two groups of cells.

**Cell-cell communication analyses**

Recently we developed a new computational tool CellChat to systematically infer and analyze intracellular communication from scRNA-seq data. CellChat infers the biologically significant cell-cell communication by assigning each interaction with a probability value (i.e., interaction score or weight) and peforming a permutation test. CellChat models the probability of cell-cell communication by integrating gene expression with prior known knowledge of the interactions between signaling ligands, receptors and their cofactors including soluble agonists and antagonists, as well as co-stimulatory and co-inhibitory membrane-bound receptors. The intercellular communication networks for the nonlesional and lesional skin were separately inferred and then jointly analyzed using CellChat (version 1.1.0). The average expression of signaling genes per cell cluster was computed using the truncated mean, where 10% of expression levels were trimmed from each end of data. Since CellChat infers intracellular communications based on cell clusters, the interactions associated with cell clusters with very few cells were potentially artifacts. We thus filtered out the inferred interactions associated with stressed keratinocyte population in nonlesional skin because of the extremely low percent of stressed keratinocytes compared to other keratinocytes in nonlesional skin (fig. 3d).

**Pseudotime and trajectory analysis**

The PHATE dimensional reduction of keratinocytes from all samples was performed by taking the shared low dimensional space obtained by scMC as an input. The parameters used in PHATE on the data are as follows: *npca = 30, t = 3*. When inferring pseudotemporal trajectory of keratinocytes, the PHATE space was used the reduced dimensional space in Monocle 3(*36*). A principal graph is learnt by learn_graph function with the parameters: *minimal_branch_len = 5*, *rann.k = 18* and *Euclidean_distance_ratio = 2*. Pseudotime values of cells were obtained once cells were ordered based on the learnt graph. In addition, we also inferred the possible transitions between different cell subpopulations using PAGA by using the PHATE space as a reduced dimensional space.

**RNA velocity analysis**

RNA velocity was calculated based on the spliced and unspliced counts as previously reported (*38*), and cells that were present in the pseudotemporal trajectory analysis were used for the analysis. We used the python implementation “scvelo” with PHATE space as an input.

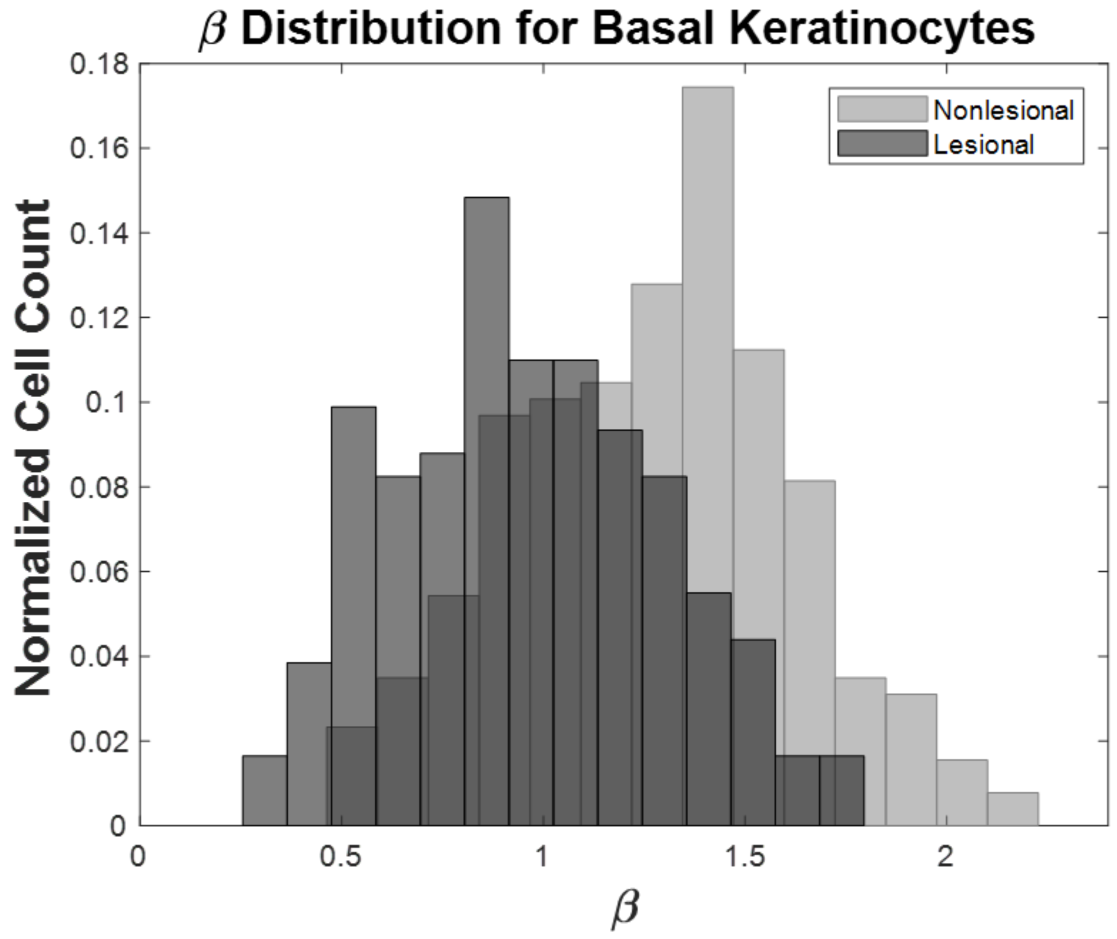


**Fig. S1. Mitochondrial Clustering Distributions for Basal Keratinocytes**. Mitochondrial clustering values were calculated for individually segmented cells from basal optical sections of vitiligo patients. The distributions were acquired by analyzing 182 cells from lesional regions and 258 cells from non-lesional regions. Counts were normalized to the corresponding cell totals.

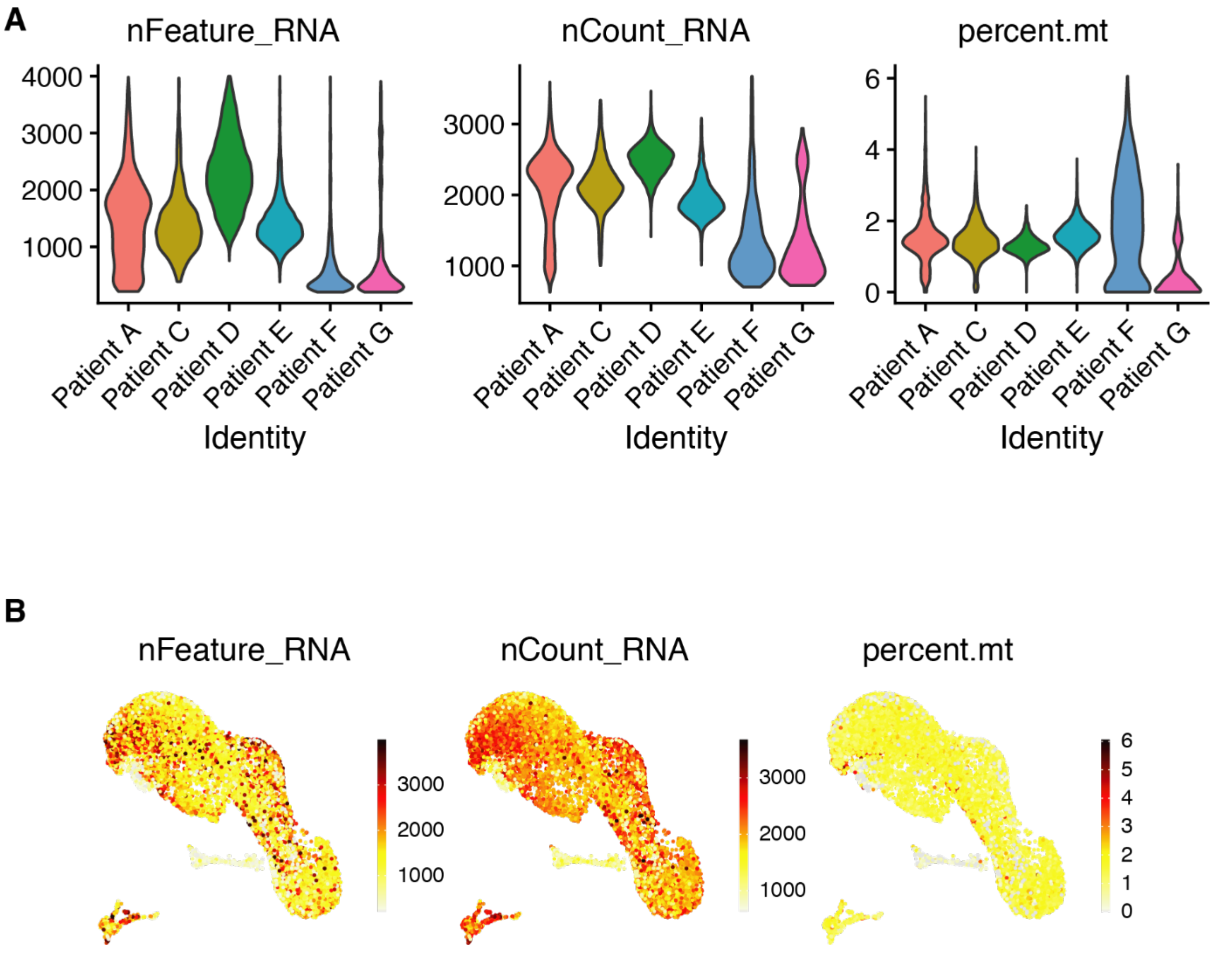


**Fig S2. Quality control metrics of the scRNA-seq data.** A) Violin plots of number of expressed genes (nFeature_RNA), number of detected counts (nCount_RNA) and percentage of mitochondrial genes (percent.mt) across all patients.   B) Overlay the quality control metrics to the UMAP plot of integrative space.

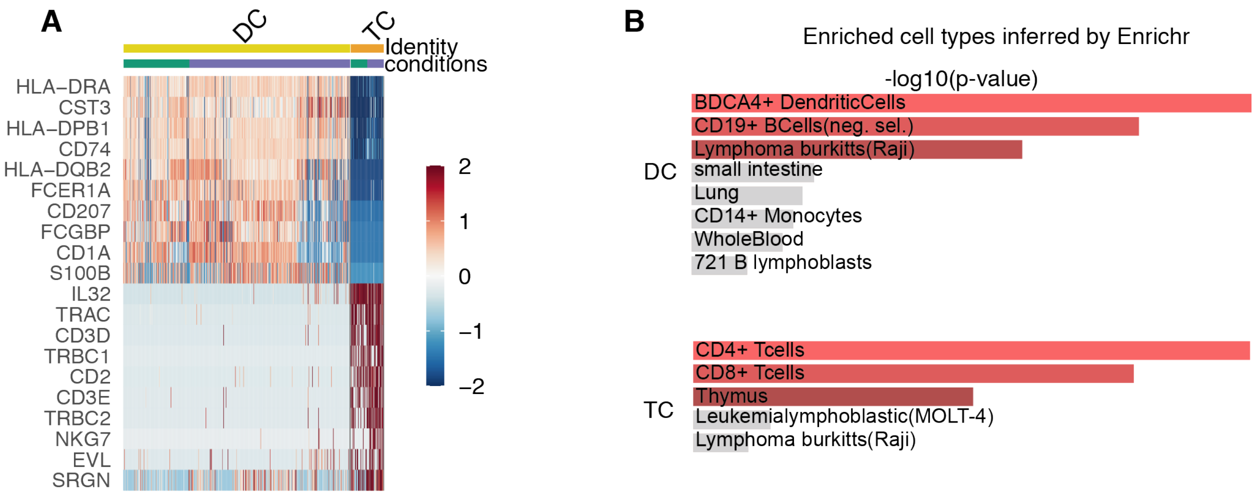


**Fig S3. The difference between DC and TC.** A) Heatmap of top 10 differentially expressed genes between DC and TC. B) Barplot of enrichment scores (-log10(p-value)) of enriched human cell types inferred by Enrichr (https://maayanlab.cloud/Enrichr/).

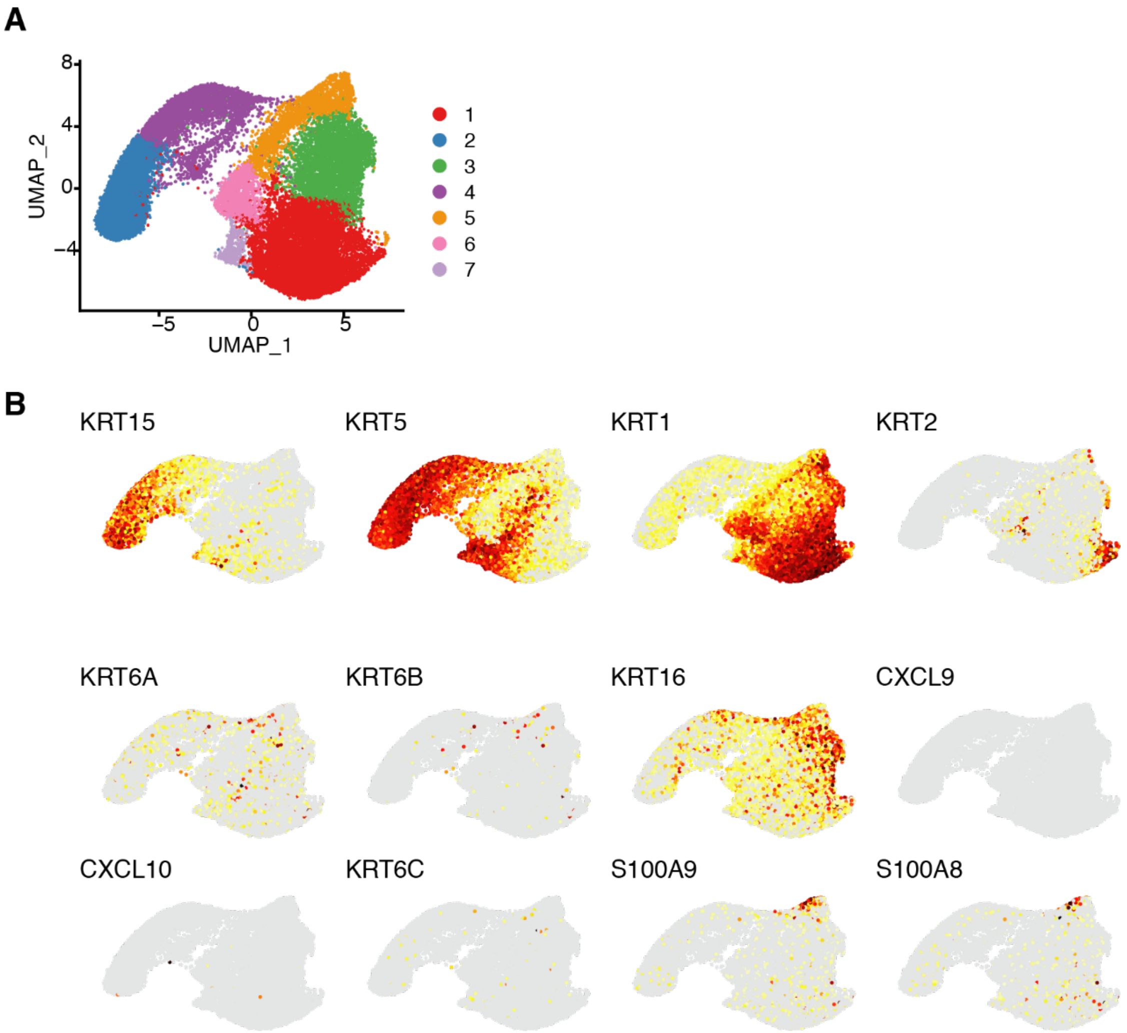


**Fig S4. Analysis results of scRNA-seq data of normal skin.** A) UMAP plot of the data with cell clusters labeled. B) Feature plots showing expression of the stressed keratinocytes markers in the UMAP plot.

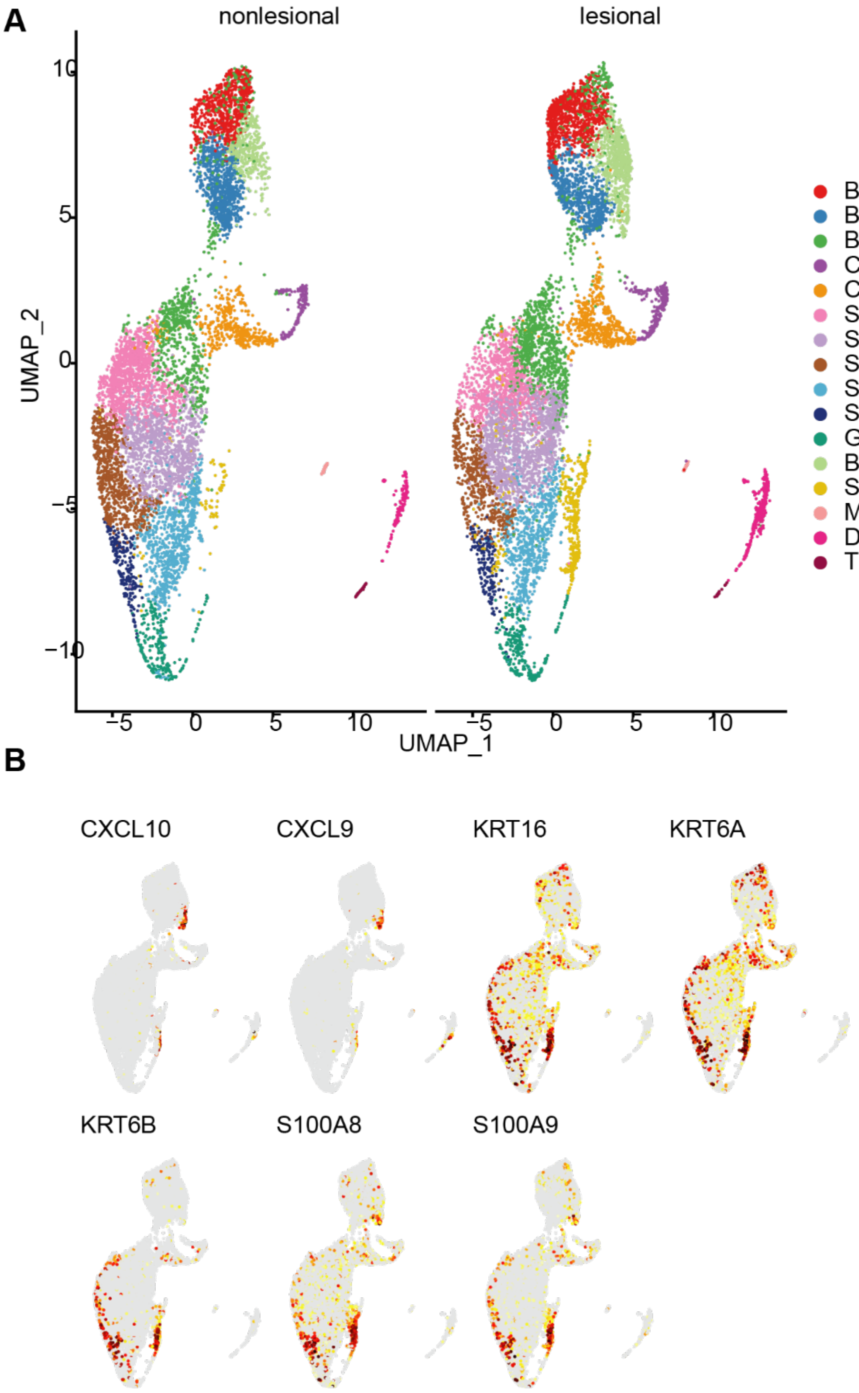


**Fig S5. Analysis results of scRNA-seq data of all patients by Seurat.** A) UMAP plot of the combined data of all patients with cell clusters labeled in both lesional (left) and nonlesional skin (right). B) Feature plots showing expression of the stressed keratinocytes markers in the UMAP plot.

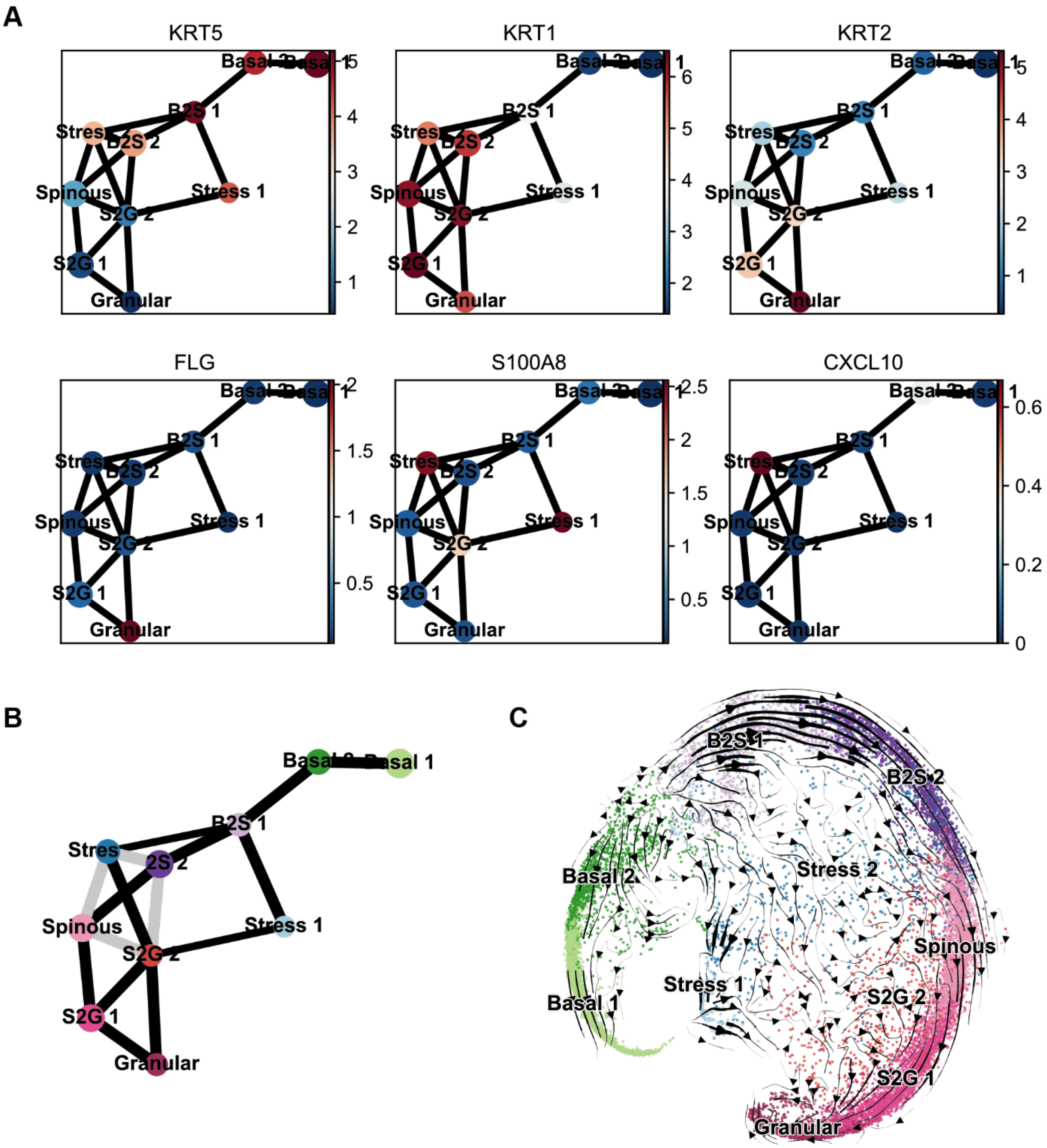


**Fig S6. Pseudotime analysis results of scRNA-seq data of all patients.** A) Marker genes' expression levels change across PAPG graph. B) PAPG graph of the PHATE space. C) RNA velocity across PHATE space.

**Table S1. Clinical Characteristics of Stable Vitiligo Patients for MPM Imaging and RNAscope**

| Patient ID | Age | Sex | Imaging Location | Disease Status |
|---|---|---|---|---|
| 1 | 61 | M | Bicep | Stable vitiligo Generalized |
| 2 | 34 | M | Hand | Stable vitiligo Acrofacial |
| 3 | 73 | F | Neck/Back | Stable vitiligo Generalized |
| 4 | 45 | F | Hand | Stable vitiligo Acrofacial |
| 5 | 74 | M | Hand | Stable vitiligo Acrofacial |
| 6 | 58 | M | Leg | Stable vitiligo Generalized |
| 7 | 36 | M | Leg | Stable vitiligo Generalized |
| 8 | 50 | F | Face | Stable vitiligo Acrofacial |
| 9 | 72 | F | Hand | Stable vitiligo Acrofacial |
| 10* | 39 | F | Leg | Stable vitiligo Generalized |
| 11* | 20 | F | Ankle | Stable vitiligo Acrofacial |
| 12* | 36 | M | Leg | Stable vitiligo Generalized |

*Denotes patients who underwent punch grafting procedure and samples were used for RNAscope

**Table S2. Clinical Characteristics of Stable Vitiligo Patients for scRNA-seq.** The number of cells after quality control are shown.

| Patient ID | Age | Sex | Areas of Involvement | Disease Status | Nonlesional cell count | Lesional cell count |
|---|---|---|---|---|---|---|
| A | 69 | F | Face, hands | Acrofacial | 604 | 1109 |
| B | 56 | F | Face, neck, back, leg | Generalized | * | * |
| C | 38 | M | Feet and legs | Generalized | 1235 | 798 |
| D | 30 | F | Back | Focal | 2729 | 2568 |
| E | 28 | M | Face, hands | Acrofacial | 2613 | 3886 |
| F | 37 | F | Legs | Generalized | 747 | 340 |
| G | 67 | F | Trunk | Focal | 553 | * |

*Insufficient viable cells and not included in analysis
